# Spatial resolution of transcriptomic plasticity states underpinning lethal morphologies in lung adenocarcinoma

**DOI:** 10.1101/2024.06.11.598228

**Authors:** Hannah L Williams, Nicolas Poulain, Ian Powley, Silvia Martinelli, Robert Bielik, Holly Leslie, Colin Nixon, Claire R Wilson, Marco Sereno, Zhangyi He, Leah Officer-Jones, Fiona Ballantyne, Rachel Pennie, Colin S Wood, David Y Lewis, Nigel B Jamieson, John Le Quesne

**Author notes:** **Corresponding author**, John Le Quesne: Garscube Estate, Switchback Road, Bearsden, Glasgow, G61 1BD. Contributed equally (Hannah L Williams, Nicolas Poulain). **Author contributions** HLW – conceptualization, data curation, formal analysis, investigation, visualization, writing-original draft, writing-review & editing. NP – data curation, formal analysis, investigation, visualization, writing-original draft, writing-review & editing. IP - data curation, investigation, formal analysis, validation, writing-review & editing. SM - data curation, investigation, validation. RB - data curation, investigation, validation. HL - data curation, investigation, validation. CN - data curation, investigation, validation. CW – data curation. MS – data curation. ZH – software, validation. LOJ - data curation, investigation, validation. FB - data curation, investigation, validation. RP - data curation, investigation, validation. CSW - data curation, investigation, formal analysis, software. DL – conceptualization, validation. NBJ – conceptualization, data curation, resources, supervision. JLQ – conceptualization, funding acquisition, supervision, writing-original draft, writing-review & editing.

## Abstract

Adenocarcinoma of the lung (LUAD) is a common and highly lethal disease. Clinical grading of disease strongly predicts recurrence and survival after surgery and is determined by morphological assessment of histological growth patterns in resected tumours. The molecular basis of growth pattern is poorly understood at present, as are the mechanisms linking growth pattern to recurrence and death.

Interestingly, the two archetypal lethal morphologies, solid and micropapillary patterns, are characterised by their biphasic appearance. Both have an epithelial fraction which is in direct stromal contact, and a fraction which is not. This morphological variance seems likely to represent plasticity, and to be causally linked to mechanisms of virulence.

To investigate the gene expression changes related to growth pattern both intra- and intertumoral, we applied spatial transcriptomics (Nanostring GeoMx DSP) to tissue microarray specimens of primary resected human lung adenocarcinoma. Using a variety of region-of-interest (ROI) selection strategies, we sampled 160 pure epithelial ROIs across 7 distinct morphological features of LUAD from 51 patients. Analyses of gene expression reveal fundamental trajectories connecting growth patterns, and crucial modes of plasticity which underly high-risk morphologies. These modes suggest mechanisms for the origins of growth pattern and mechanisms of virulence.

Our work highlights dramatic divergence in gene expression programmes between highly lethal but morphologically diverse modes of tumour growth. Furthermore, it provides an explanation for how microscopically localised hypoxia in the primary tumour helps to establish and maintain survival strategies which ultimately determine morphology-specific mechanisms of tumour metastasis, suggesting new therapeutic vulnerabilities.

## Introduction

Lung adenocarcinoma (LUAD) is the second most common malignancy in the world^1^ and one of the most lethal, with only 10% of patients surviving longer than 10 years. Histologically, LUAD tumours are composed of a diverse range of growth patterns (lepidic, acinar, cribriform, solid, papillary and micropapillary) with multiple growth patterns occurring within the same tumour. Tumours demonstrating ≥20% solid, micropapillary or complex glandular components are considered high-grade^2^ and are highly virulent, recurring more commonly post-resection and with worse overall survival^3,4^. Furthermore, the two archetypal high-grade growth morphologies, solid and micropapillary patterns, are morphologically biphasic. In solid areas, only the cells at the edge of tumour nests contact stroma, while in micropapillary tumours, only the cells not forming micropapillae, i.e., the sessile sheets from which the micropapillae emerge, are stroma-contacting. We set out to discover the molecular underpinnings of growth pattern, to see if the biphasic nature of high-grade growth patterns reflects underlying epithelial plasticity, and to link these molecular features to mechanisms of lethality.

To date, efforts to elucidate the molecular mechanisms defining specific growth patterns have been met with only partial success. There are associations between worsening degrees of grade/dedifferentiation and mutational burden/genomic complexity^5,6^. The advent of spatial transcriptomics methods has started to reveal gene expression differences between growth patterns^7,8^, but the observed molecular states have proven difficult to interpret in terms of their relationship to morphology and virulence. This is in part due to methodological limitations such as the use of bulk RNA sequencing to resolve growth pattern specific signatures in samples where mixed growth patterns are common, and signal noise from the tumour microenvironment is high. Thus, even epithelial targeted sequencing (through for example, laser capture microdissection) analysis often returns a “mixed” readout representative of more than one growth pattern^5^.

The recent development and commercial availability of spatially resolved transcriptional sequencing methods such as the Nanostring GeoMx Digital Spatial Profiler (DSP)^9^ now enables the assessment of growth pattern specific transcriptional programs with extremely high resolution and sample purity. Few studies have utilised this technology to resolve growth pattern identity and those which have still fail to isolate unique growth patterns in analysis^10^.

To address this, we present our study using spatially resolved transcriptomic sequencing to elucidate growth pattern-specific biology and to further investigate intra-growth pattern heterogeneity of high-risk growth patterns. This was performed in a cohort of formalin fixed paraffin embedded tissue microarrays comprising primary lung adenocarcinoma from 51 unique patient samples. Through the application of bespoke region of interest (ROI) selection strategies to separate regions within tumour epithelium and subsequent analyses of gene expression data, we reveal unique molecular programs related to growth patterns, as well as intra-tumoural phenotypic switching. Our work highlights dramatic divergence in phenotype between the two archetypal highly lethal tumour morphologies and provides an explanation for how microscopically localised hypoxia primes groups of cells for survival and metastatic spread by two distinct naturally selected strategies, suggesting new therapeutic vulnerabilities.

## Methods

### Cohort

The cohort (LATTICe-A: Leicester Archival Thoracic Tumour Investigatory Cohort – Adenocarcnoma) used for this study comprises a 994-patient retrospective cohort of resected lung adenocarcinomas with comprehensive clinicopathological annotation including survival information and has been previously described ^11,12^. A subset of this cohort (Table.S1) (four tissue microarray blocks) was used for the spatial transcriptomics experiments described below. This comprised 51 unique patients from which adenocarcinoma samples were profiled. The LATTICe cohort was originally constructed under REC 14/EM/1159 (East Midlands REC), and the ongoing management of the resource by the Greater Glasgow and Clyde Biorepository was approved under an amendment granted by the Leicester South REC. Ongoing use of the collection is now managed under REC 16/WS/0207.

### Spatial transcriptomics using Nanostring GeoMx DSP

The basic summary of the DSP experiment consisted of primary mask visualisation, ROI and AOI segmentation, probe retrieval and sequencing and downstream data analysis (Fig.S1).

Sample preparation for Nanostring GeoMx DSP: Detailed slide preparation has been previously described^13^. Primary visualization antibodies, PanCK - epithelium (Cy5) and Syto13 - nuclei (FITC) were used to aid ROI selection. Epithelium was selected based upon PanCK^+^ expression. Each TMA was cut onto a separate slide and hybridized with probes from the Whole Transcriptome Atlas (WTA) targeting 18,000 protein coding genes.

Region of interest and area of illumination selections: Following hybridization, slides were scanned on the GeoMx DSP instrument and regions of interest (ROIs) and areas of illumination (AOIs) selected using the GeoMx DSP analysis suite. Cores containing growth patterns (normal, in-situ, acinar, cribriform, and papillary were identified and confirmed using H&E by a pathologist. Geometric segmentation tool was used to select ROIs for these respective morphologies. Epithelial AOIs were selected within each ROI using positive panCK selection. For cores with micropapillary growth patterns we used the polygon segmentation tool to first isolate floret and sessile ROIs and subsequently select epithelial AOIs within each of these respective ROIs using positive panCK selection. For cores with solid growth patterns, we first selected solid “islands” using the polygon segmentation tool and exported each ROI to ImageJ. In ImageJ we applied a modified macro (Nanostring) to grow contours from the solid island edge in 20µm increments to generate a maximum of three compartments (edge, intermediate and center). A mask was generated from each compartment and imported back into the GeoMx DSP interface, aligned with the original image.

Probe retrieval and sequencing: Upon completion of AOI selection, photocleavable oligonucleotide probs were exposed to UV-light, cleaved and the subsequent aspirate was collected into a 96-well DSP collection plate. Following rehydration with DEPC-treat water, the probes were added to the corresponding well of a new 96-well PCR plate which contained GeoMx Seq Code primers and PCR master mix. Full details of library preparation and sequencing have been previously described^13^.

### Data analysis of spatial transcriptomic data

AOI and target filtering: Following initial QC steps including biological probe assessment, AOIs with less than 250 genes above limits of quantitation (LOQ: the limit of quantitation (LOQ) for each AOI is calculated as 2 geometric standard deviations above the geometric mean of the negative probes. This is calculated after the exclusion of outlier probes) were excluded from further analysis. Targets with less than 1.5% of AOIs (equivalent of 3 segments from total sampled) above LOQ were also excluded from downstream analysis (subsequent genes n = 14,967 genes). Sixteen ROIs from 6 patients were removed from further analysis due to classification as mucinous cases (173/189 ROIs). Finally, further AOI filtering to exclude AOIs with <60% aligned reads (the percentage aligned reads were calculated, aligned reads/raw reads*100), 13 ROIs failed to meet these criteria and were thus removed from further analysis, n AOIs = 160/189) (Fig.S1).

Data preparation for differential gene expression, morphology: AOIs from cores representing solid and micropapillary growth patterns were collected from spatially distinct regions: Solid = three contour AOIs per solid “island” and Micropapillary = AOIs from floret and sessile structures. Comparisons were performed between the following morphological classifications:

1. Macro growth patterns (dataset 1): normal (*n=5)*, in-situ (*n=5)*, acinar (*n=9)*, cribriform (*n=12)*, solid (*n=12)*, papillary (*n=8)*, micropapillary (*n=12)*: total = 63.
2. High risk solid growth pattern intra-ROI sampling (dataset 2): solid-centre (n AOIs = 12), solid-intermediate (n AOIs = 11), solid-edge (n AOIs = 10): total 12.
3. High risk micropapillary growth pattern intra-ROI sampling (dataset 3): micropapillary-floret (n AOIs = 11), micropapillary-sessile (n AOIs = 11): total 11.

In the instance of multiple cores of the same morphology from the same patient gene counts were averaged so that one morphology per patient was represented in dataset 1, this method was also applied to dataset 2 solid growth patterns with contoured sampling and dataset 3 micropapillary growth patterns split into sessile and floret AOIs.

We applied principal component analysis to examine potential batch effects from deriving data from different TMA blocks (Fig.S2A) or sequencing plates (Fig.S2B). While we did observe marginal data segregation by TMA blocks the growth pattern distribution was not equal across each block thus, we could not deduce whether this was true batch effect or underlying growth pattern specific biology. For these reasons we did not perform batch adjustment on the dataset.

Differential gene expression, gene set variation analysis and pathway enrichment analysis: Differential gene expression was performed using R package DESeq2^14^. Non-normalised read counts were input into the DESeq2 pipeline which performs median of ratios normalization. For differential gene expression analysis, independent filtering was applied to remove genes with low counts and Cooks cut-off, to remove outliers prior to further downstream analysis.

Gene set variation analysis (GSVA) was performed using the GSVA^15^ R package. Enrichment scores per sample for the Hallmarks gene-sets were averaged across growth patterns. This average value per growth pattern was further used to derive enrichment scores for more generalised hallmarks (apoptosis, cellular component, cellular stress, development, DNA damage, EMT, hypoxia, immune, metabolism, proliferation and, signalling). 50 hallmarks pathways were ranked for each growth pattern and each rank normalised by the maximum rank (50) to scale the values between 0-1. In the instance of when multiple gene-sets were comprised in a single generalised hallmark e.g., “proliferation” contained 6 hallmarks pathways, normalised ranks for these gene-sets were averaged to derive a single score for each generalised hallmark per growth pattern. To enable comparison of the enrichment of our derived generalised hallmarks for each growth pattern, these scores were finally normalised within each growth pattern, diving each value by the maximum score for that specific growth pattern such that a value of 1 would represent the highest degree of enrichment of a single generalised hallmark.

Pathway analysis was performed using the R package clusterprofiler^16,17^. To examine enrichment of specific biological domains in gene set enrichment analysis (GSEA) we used Hallmarks^18^, Reactome^19^ and GO^20,21^ pathway gene sets. Further investigation of enriched gene sets was performed by assessing the top 10 gene overlaps between each enriched Hallmarks pathway and Reactome and GO gene sets respectively using the interface available on the MSigDB online database. Visualisation was performed using R packages clusterprofiler and ggplot2^22^. Benjamini-Hochberg was performed to account for multiple testing and an adjusted p-value of 0.05 or 0.01 (specified) was applied representing either a 5% or 1% false discovery rate.

Trajectory analysis: The dataset was first restricted to a single core per case, using an average of the ROIs for each growth pattern per core. The 10% most biologically informative genes were selected based on a mean-variance trend in gene expression, using the Bioconductor “scran” package. Three clusters were identified by application of a Leiden clustering algorithm to the principal component analysis (PCA) dimensional reduction plot based upon the selected genes; the three clusters are enriched for in situ, solid and micropapillary growth respectively. The cluster containing all in situ samples was designated as the origin, and unsupervised application of the “Slingshot” algorithm identified two trajectories, i.e. the two clusters containing invasive patterns are connected by separate pathways and are not in series. Samples excluded from the computation of trajectory were then reintroduced by projecting them in the PCA space. Pseudotime, i.e. distance to the origin of the projections of samples upon inferred trajectories, was then calculated for all samples. The same method was applied to a publicly available dataset GSE58772^23^, obtained by RNA sequencing of laser capture microdissected epithelial cells from fresh-frozen tumour tissue. To represent pseudotime in a way that could be exported to other datasets, we defined scores similarly to the Tavernari lepidic to solid score^24^. Scores of transitions from lepidic to solid (L2S) and from lepidic to micropapillary (L2M) were calculated as the weighted sum of z-normalised expression of the most differentially expressed genes between lepidic and the respective high-grade pattern (FDR<0.05 and log2(FC)>2).

### Tempo-seq

TempO-seq targeted sequencing-based RNA expression sequencing was performed. 713 non-mucinous LUAD cores from 23 FFPE TMA slides (Table.S2) were scraped and lysed using the TempO-Seq 2X Lysis Buffer. The samples were analysed using the Human Whole Transcriptome v2.1 panel with standard attenuators. Quality control of resulting data was performed, including density, clusters purity factor, quantity, and quality of reads. TempO-Seq sequence data were pre-processed using the Tempo-Seq-R^TM^ software package, reads were aligned using the STAR^25^ algorithm to a pseudo-transcriptome corresponding to the gene panel used in the assay. Gene isoforms were merged, and genes detected in less than a third of samples were removed. Our TempO-seq gene expression dataset was then normalised using upper-quartile method and a log2 transformation. The level of the L2S and L2M score was calculated using z-normalised expression of genes included in signatures. Score values were then compared between predominant histological pattern using anova and each pattern against all other samples using Student test. Survival analysis were performed using log-rank regression as implemented in the “Survival” R package.

### Multiplex immunofluorescence

The first mIF assay was used to detect Human Glucose Transp. 1(Glut-1 ) , Pan-Cytokeratin (AE1/AE3), and Fatty Acid Synthase (FASN). Sections were cut at 4um thickness on TOMO slides. The slides were baked at 60°C for 1 hour, the rest of the process was fully automated on the Ventana Discovery Ultra (Roche Tissue Diagnostics, RUO Discovery Universal V21.00.0019). Slides were dewaxed and antigen retrieval was performed using Discovery Cell Conditioning 1 (Roche Tissue Diagnostic, ref. 06414575001) for 32 minutes at 95°C. FASN (C20G5) (Cell Signalling Technology, ref.3180) was applied at 1:25 for 60 minutes at 37°C, followed by DISCOVERY OmniMap anti-Rb HRP secondary antibody (Roche Tissue Diagnostics, ref. 05269679001) for 24 minutes, detected by the fluorophore opal 520 at 1:200 (Akoya, ref. FP1487001KT) for 8 minutes. A denaturation step was performed using Cell Conditioning 2 (Roche Tissue Diagnostics, ref. 05279798001). Glut-1 (2B scientist, ref GT12-A) was applied at 1:150 for 32 minutes at 37°C, followed by DISCOVERY OmniMap anti-Rb HRP secondary antibody (Roche Tissue Diagnostics, ref. 05269679001) for 12 minutes, detected by the fluorophore opal 690 at 1:100 (Akoya, ref. FP1487001KT) for 8 minutes. A denaturation step was performed using Cell Conditioning 2 (Roche Tissue Diagnostics, ref. 05279798001). AE1/AE3 (Leica Biosystems, ref. AE1/AE3-601-L-CE) was applied at 1:250 for 28 minutes at 37°C, followed by DISCOVERY OmniMap anti-Ms HRP secondary antibody (Roche Tissue Diagnostics, ref. 05269652001) for 12 minutes, detected by the fluorophore opal 620 (Akoya, ref. FP1495001KT) at 1:50 for 8 minutes. DAPI (Roche Tissue Diagnostics, ref. 05268826001) was used as a nuclear counterstain. We then manually mounted the slides using Diamond prolong (Thermo Fisher, ref. P36970). Whole slide images were collected at 10x magnification using the PhenoImager multispectral slide scanner (V1.0.13, Akoya Biosciences). A TMA map was applied using Phenochart (v1.1.0, Akoya Biosciences) and individual core images acquired at 20x magnification. Single stained images of human lung and an unstained section image were collected using the PhenoImager to spectrally unmix the core images using Inform (version 2.5.1., Akoya Biosciences). Component files were imported to Qu-Path for image analysis. All antibodies were validated on the manufacturers recommended tissue control using the clinical gold standard (chromogenic DAB) assay to ensure specificity and sensitivity prior to fluorescent optimisation. Single fluorescence assays were used to define the optimal dilution factor of the fluorophore for the multiplex assay. A negative control multiplex assay was designed replacing the test antibodies with CONFIRM Negative Control Rabbit Ig, (Roche tissue Diagnostics, ref.760-1029) and Negative Control Mouse Monoclonal Antibody (MOPC-211), (Roche tissue Diagnostics, ref. 760-2014) and used on each multiplex staining run.

A second mIF assay was used to quantify proliferation. The slides were baked at 60°C for 1 hour, the rest of the process was fully automated on the Ventana Discovery Ultra (Roche Tissue Diagnostics, RUO Discovery Universal V21.00.0019). Slides were dewaxed and antigen retrieval was performed using Discovery Cell Conditioning 1 (Roche Tissue Diagnostic, ref. 06414575001) for 32 minutes at 95°C.

The assay consisted of the following antibodies and corresponding dilution factor: CK AE1/AE3 (Leica Biosystems, ref. AE1/AE3-601-L-CE) was applied at 1:250 for 28 minutes at 37°C, followed by DISCOVERY OmniMap anti-Ms HRP secondary antibody (Roche Tissue Diagnostics, ref. 05269652001) for 12 minutes, detected by the fluorophore opal 620 (Akoya, ref. FP1495001KT) at 1:100 for 8 minutes and KI-67 (Roche Tissue Diagnostic, ref. 05278384001) was applied for 20 minutes at 37°C, followed by DISCOVERY OmniMap anti-Rb HRP secondary antibody (Roche Tissue Diagnostics, ref. 05269679001) for 12 minutes, followed by TSA-DIG (Akoya Bioscience, ref. FP1501001KT ) at 1:100 for 10 minutes and detected by the fluorophore opal 780 (Akoya, ref. FP1501001KT) at 1:25 for 60 minutes.

Whole slide images were collected at 10x magnification, and individual core images were collected at 20x magnification as previously described. Core images were spectrally unmixed using InForm (v 4.2, Akoya Biosciences), and component images were imported to Visiopharm for analysis. A deep learning classifier was developed using Visiopharm (version 2021.09.2.10918) and used to detect regions of necrosis, tumour, stroma, and background tissue for subsequent cell level analysis. A further deep learning classifier was used to detect cells and generate cell level mean intensity measures.

## Results

**Two inferred trajectories to high-grade malignancy**

Lung adenocarcinomas (LUADs) are made up of diverse epithelial morphologies, histopathologically classified as in situ, acinar, cribriform, solid, micropapillary and papillary growth patterns (Fig.1A). This forms the basis of histopathological tumour grading^26^, and it is well established that tumours with a predominance of solid or micropapillary growth patterns have poor patient outcome, and this is confirmed in our cohort (Fig.1B).

**Figure 1.**
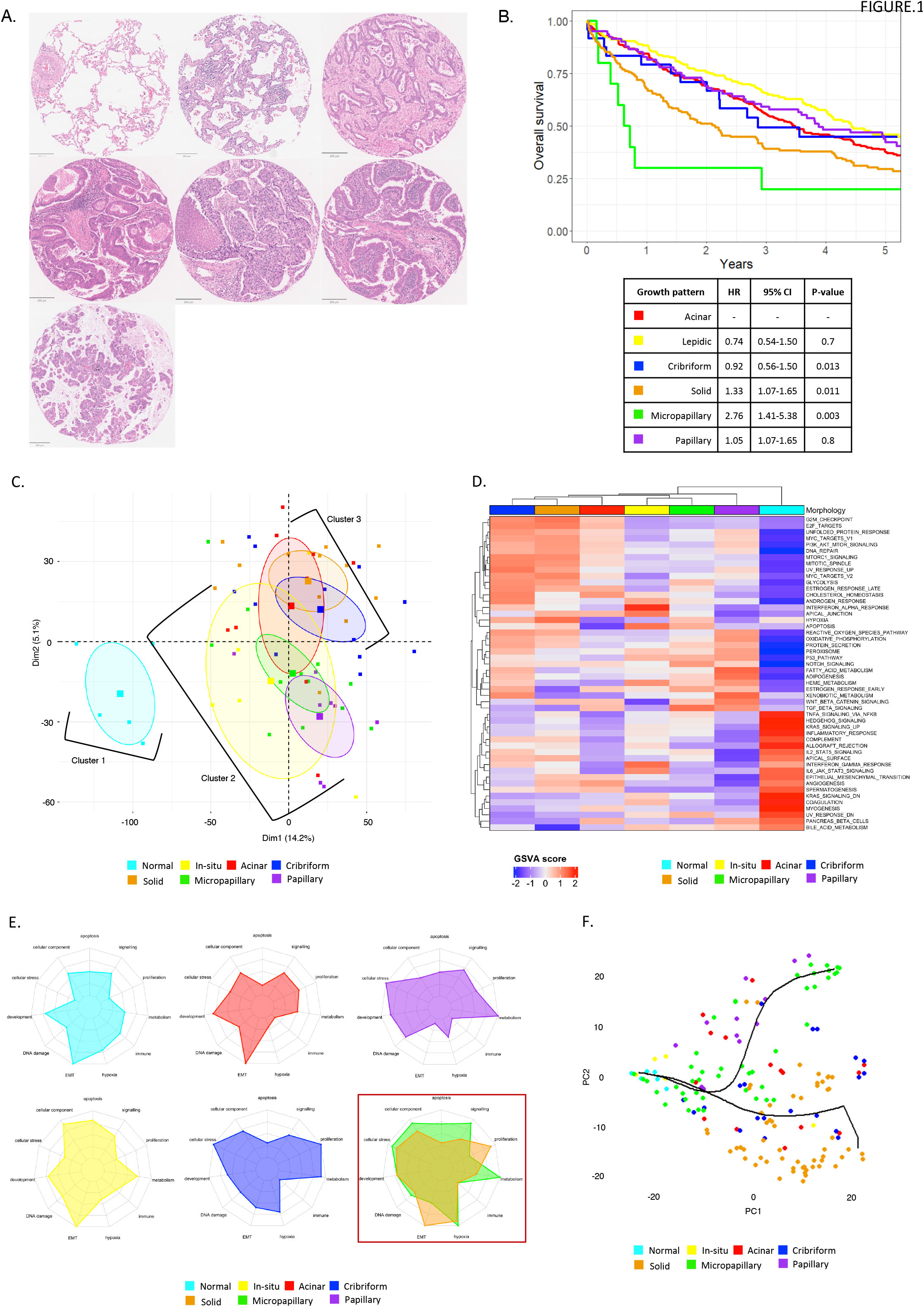
A: Haematoxylin and eosin representative images of growth patterns from cohort. Left to right: normal, in-situ, acinar, cribriform, solid, papillary and micropapillary (scale bar 200μm). B: Survival analysis of predominant histological pattern. Top panel is the Kaplan-Meier representation and bottom panel represents corresponding table of estimates of corresponding Cox regression analysis. C: Principal component plot of gene expression from dataset 1. PCA shows three groupings of morphologies: cluster 1 – normal, cluster 2 – in-situ, micropapillary and papillary and, cluster 3 – acinar, cribriform and solid growth patterns (colour key: cyan – normal, yellow – in-situ, red – acinar, blue – cribriform, orange – solid, green – micropapillary, purple – papillary). D. Averaged gene set variation score derived from per sample gene set variation analysis using Hallmarks pathway gene-set. E. Radar plots of concatenated hallmarks for each growth pattern. E: Unsupervised slingshot trajectory analysis of pure epithelial samples.

To gain global insight into the distribution of epithelial gene expression profiles of different LUAD growth patterns we derived expression data on 14,947 genes using the digital spatial profiler spatial transcriptomic platform (Nanostring GeoMX), providing global transcriptomic coverage and excellent growth pattern purity. Following data processing (Methods), we performed principal component analysis on case level gene expression. This highlighted a degree of separation between growth patterns (Fig.1C) with three main clusters of morphologies: 1. Normal epithelium, 2. in-situ/micropapillary/papillary, 3. acinar/cribriform/solid.

We next calculated the average GSVA score of the hallmarks gene ontology database (Table.S3) per growth pattern to examine growth pattern specific transcriptomic features (Fig.1D). To further resolve differences between growth patterns, we generated generalised hallmark terms (Methods) and scored each growth pattern for their specific enrichment for each term (Fig.1E). This enabled identification of clear differences between growth patterns. Normal alveolar epithelial cells demonstrated a distinct transcriptomic profile from all malignant growth patterns, consistent with relatively low proliferation, cellular stress, and metabolic activity. Acinar, cribriform, and solid growth patterns all demonstrated enrichment for proliferation and DNA damage hallmarks. Interestingly, the two archetypal high-risk morphologies, solid and micropapillary patterns, demonstrated markedly contrasting transcriptomic profiles. While both showed enrichment for hypoxia-related genes, micropapillary growth patterns lacked the enrichment of proliferative and EMT programs observed in solid growth, instead showing an enrichment in hallmarks related to metabolism and cellular structure. The extent of these differences was further confirmed from pairwise differential gene expression (Table.S4) in which solid and micropapillary regions demonstrated a large relative percentage of differentially expressed genes out of all comparisons (1.23% of total genes significantly differentially expressed, p.adj≥0.01).

To further study the relationship between growth patterns, and with the hope of inferring possible routes of tumour dedifferentiation and to further discern differences between solid and micropapillary growth, we performed an analysis which identified two distinct trajectories in mRNA expression space (Fig.1F). The origin was set as in situ/lepidic growth, based on the known biological progression of invasiveness from these lesions^27^. The trajectories share an early path but then branch towards areas dominated by cribriform and solid ROIs and by papillary and micropapillary ROIs respectively. These observations were validated in a publicly available dataset^23^ of laser micro dissected pure epithelial gene expression which lacked only cribriform histological pattern (Fig.S3A). To quantify the position along these two paths toward high-grade disease, we defined two metagenes, derived from the most differentially expressed genes between lepidic and solid (L2S) and lepidic and micropapillary (L2M) ROIs. As expected, solid growth has a significantly higher L2S score in comparison to in situ, papillary and micropapillary regions (Anova, p=2.2^e-16^), while micropapillary samples have higher L2M score. Subregions of solid growth patterns all have similar L2S values, while sessile subregions have a slightly higher L2M values than micropapillary ROIs (Fig.S3B&C).

To evaluate if these signatures retained meaning in bulk transcriptomic data, we calculated the level of expression of L2S and L2M signatures in TempO-seq data obtained from DNA extracted from 713 cases across 23 TMA sections, including the 4 TMAs upon which DSP was performed. We observed similar trends of L2S and L2M increasing towards respective predominant high-grade patterns (Fig.S2D&E), indicating that the epithelial signature predicts growth pattern progression despite stromal signal noise. To evaluate the potential clinical relevance of these scores, we assessed their association with overall survival. Both scores were strongly associated with a reduced OS and remained significant when included in a Cox model (L2S: HR 1.87, 95% CI 1.54-2.26, p<0.01, L2M: HR 1.46, 95% CI 1.16-1.85, p=0.001), showing that they encompass at least partly independent information (Fig.S2F&G).

### Solid and micropapillary

We next sought a head-to-head comparison of two lethal growth patterns (solid and micropapillary) which happen to show one of the largest transcriptomic divergences of all invasive growth patterns (surpassed only by solid vs papillary growth) as demonstrated by pairwise differential gene expression (Table.S4). We identified 120 significantly differentially expressed genes (p.adj≤0.01, Fig.2A&B, Table.S5). Eighty-six genes were significantly upregulated in solid growth patterns compared to 34 genes significantly upregulated in micropapillary. Significantly upregulated genes in solid growth patterns include several associated with cell motility and cytoskeletal remodelling e.g., *MARCKS, TPX2* and *FBLN1*. In additional several genes associated with cell cycle were also significantly upregulated e.g., *MYBL2, PCNA* and *CDK4*, as well as numerous chromatin constituents. In the case of micropapillary growth patterns, the most upregulated genes encode proteins secreted by type II alveolar cells (SCGB3A2, PIGR, MUC1), followed by genes associated with metabolic processes, including that which limit glycolysis under conditions of hypoxia (*TXNIP* and *FABP3)*. Gene set enrichment analysis (Fig.2C) further confirmed proliferation-related hallmark features enriched in solid growth pattern areas (E2F targets, MYC targets and G2M checkpoint genes) whilst micropapillary growth patterns demonstrated an enrichment in metabolic hallmarks such as fatty acid metabolism (padj<0.05).

**Figure 2.**
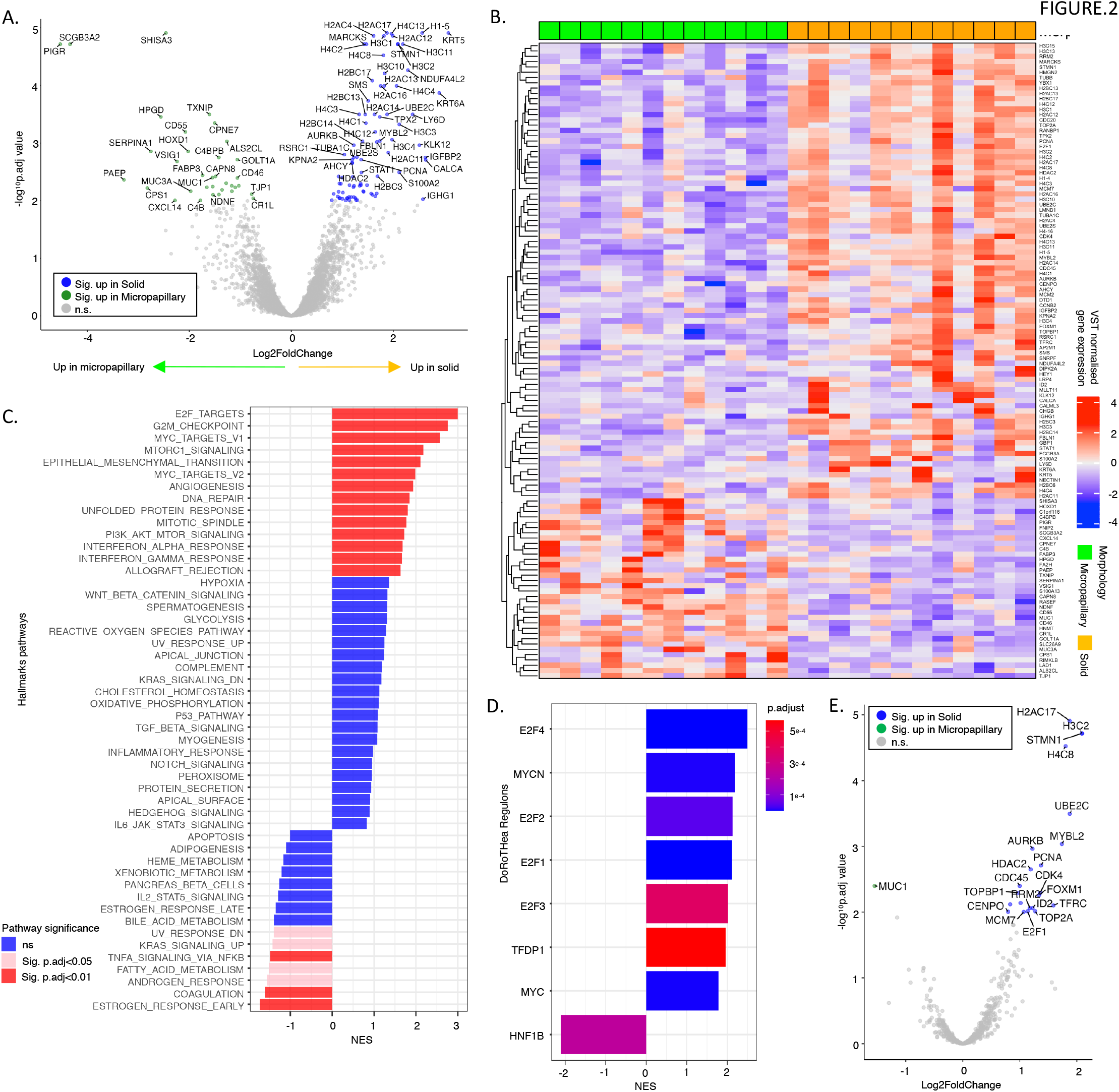
A: Volcano plot of differentially expressed genes (DEGs) for comparison of solid v. micropapillary from dataset 1 (sig values = p.adj <0.01). B: Sample wise gene expression of significantly (p.adj <0.01) DEGs for solid and micropapillary samples (red: micropapillary, blue: solid). C: Gene set enrichment analysis of solid v. micropapillary gene expression using Hallmarks gene-sets (blue: not significant, pink: p.adj ≤0.05, red: p.adj ≤0.01). D: Network analysis using DoRoTHea regulon gene-sets. E. Volano plot of significantly DEGs comprised in solid growth pattern enriched DoRoTHea regulons.

To gain further insight into potential master transcriptional regulation of the differences observed we performed gene regulatory network (GRN) analysis using the DoRothEA GRN^28,29^ regulon sets to infer transcription factor (TF) activity. Six regulons were significantly enriched in solid growth patterns (Fig.2D) including two MYC regulons and 4 members of the E2F family of transcription factors. Investigation into the differential gene expression of specific components of each regulon (Fig.2E) demonstrated significant upregulation associated with cell cycle, cellular proliferation, and cellular invasion e.g., CDK4 (p.adj=0.005), FOXM1 (p.adj=0.005) and PCNA (p.adj=0.001) implicating the E2F family of transcription factors as having an important role in shaping the solid phenotype. No regulons were identified as significantly enriched in micropapillary growth pattern.

### Solid growth pattern islands are molecularly biphasic

Having identified differences in gene expression between entire epithelial regions with these two high-risk morphologies, we next sought evidence of intra-tumoral epithelial plasticity related to their diverse internal microenvironments. We started by comparing the stroma-contacting (peripheral) vs stroma-isolated (central) regions of solid growth pattern (Fig.3A). To assess the transcriptomic profiles of these in a spatially resolved manner we segmented solid tumour islands into three contouring AOIs: edge (0-20µm from stromal interface), intermediate (20-40µm from stromal interface), and centre (Fig.3B). After averaging gene expression across multiple solid region segments from the same patient, we analysed solid regions from 11 patients (33 AOIs: 10 edge, 11 intermediate, 12 centre).

**Figure 3.**
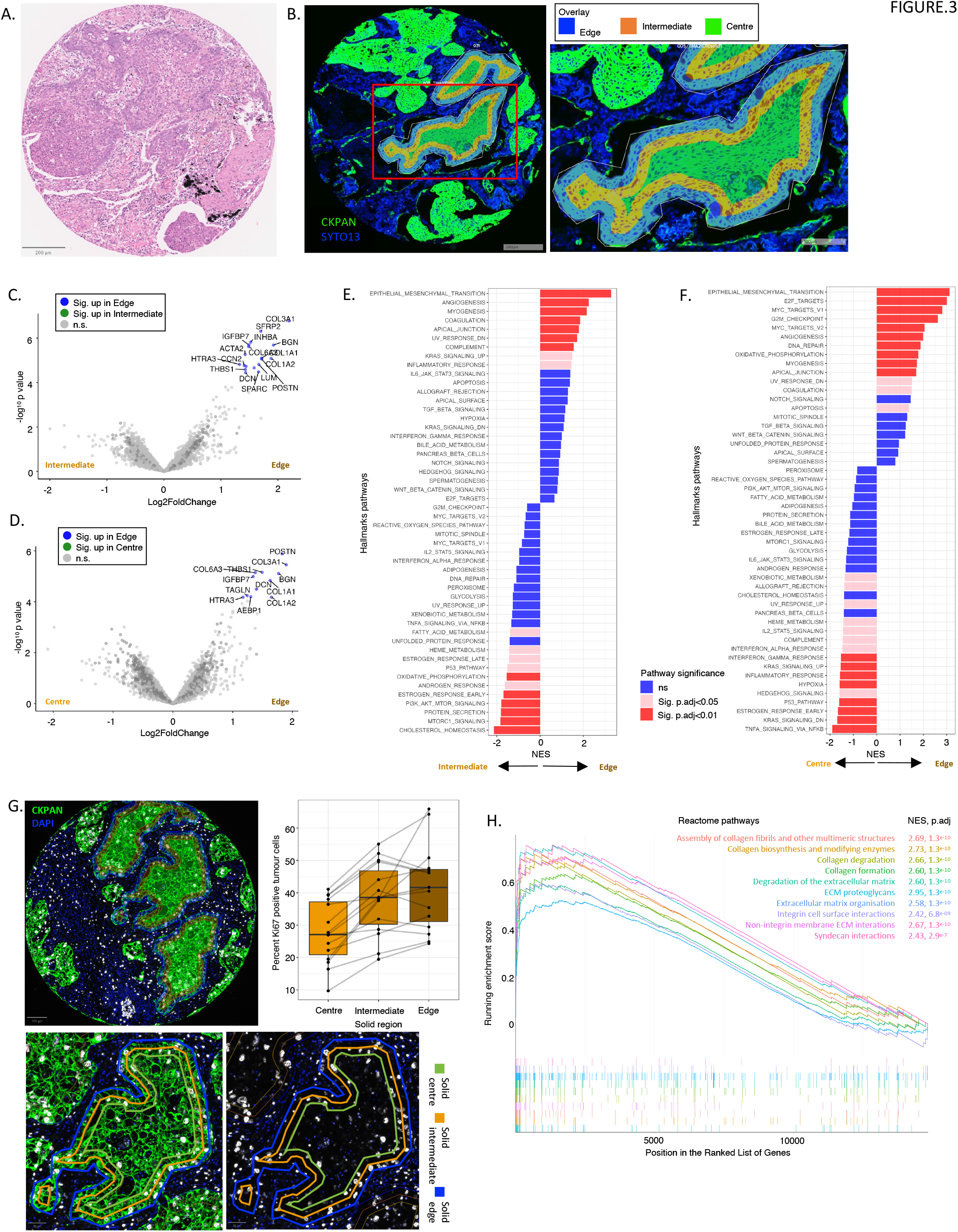
A: Haematoxylin and eosin stained representative image of solid growth pattern (scale bar 200μm). B: Example segment selection on Digital Spatial Profiler (Nanostring) to create three contours per solid island (left image scale bar 200μm, right image scale bar scale bar 100μm). C: Volcano plot of significantly differentially expressed genes (DEG) in solid_edge v. solid_intermediate contours (p.adj ≤0.05). D: Volcano plot of significantly differentially expressed genes (DEG) in solid_edge v. solid_centre contours (p.adj ≤0.05). E: Gene set enrichment analysis (GSEA) of solid_edge v. solid_intermediate gene expression using Hallmarks gene-sets (blue: not significant, pink: p.adj ≤0.05, red: p.adj ≤0.01). F: GSEA of solid_edge v. solid_centre gene expression using Hallmarks gene-sets. G: Representative multiplex immunofluorescent images of Ki67 expression (surrogate of proliferation) in solid growth. Increase in Ki67 expression observed in edge and intermediate contours in comparison to centre (top left: scale bar 100μm, bottom left and right: scale bar 50μm). Boxplot demonstrates percent Ki67 positive cells in each contour per sample. Significant difference between intermediate and edge with center segments (Kruskal-Wallis: p=0.0081). H: Edge plots of significantly enriched Reactome pathways comprising overlapping genes with Hallmarks EMT pathway in solid_edge v. solid_centre. (s p-value < 0.1, * p-value < 0.05, ** p-value < 0.01, *** p-value < 1e-3).

Differential gene expression analysis of contouring regions demonstrated that peripheral/edge contours expressed 18 and 12 upregulated genes compared to intermediate (Fig.3C) and centre (Fig.3D) respectively (p.adj≤0.05), with significant overlap. No significantly differentially expressed genes were identified between inner and centre (data not shown). Overall, peripherally located tumour cells exhibited upregulation of extracellular matrix components (e.g., collagens, decorin, thrombospondin). Further analysis by GSEA (Fig.3E&F) demonstrated significant enrichment of hallmarks related to EMT in the edge zone compared to both other zones, and a further marked relative upregulation of proliferation-related hallmarks compared to centrally situated cells. This was validated through immunolabelling of Ki67 positive cells by quantitative immunofluorescence (qIF) on the same tissue cores that were sequenced. Again, solid islands were segmented into three regions and the percentage Ki67 positive tumour cells within each region was quantified. This confirmed observations at RNA level that proliferation is significantly increased in edge and intermediate compared to the solid centre (Kruskal-Wallis test: p=0.0081, Mann-Whitney test: edge v. centre: p=0.0032, intermediate v. centre: p=0.0032) but not significantly different between intermediate and edge regions (p=0.59) (Fig.3G).

To gain a more granular insight into the flavour of biological processes associated with the genes involved in the EMT hallmarks we calculated the top 10 Reactome pathways which comprised significantly overlapping genes with the hallmarks EMT geneset. Specifically, pathways associated with ECM remodelling (“Collagen degradation”, “degradation of the ECM”, “ECM organisation”) and ECM engagement for cellular motility (“integrin cell surface interactions”, “non-integrin membrane ECM interactions”) were significantly enriched in both edge vs intermediate and edge vs centre (Fig.3H, Table.S6) comparisons. Elevation of these pathways at the stromal interface indicates a distinct locally aggressive and proliferative cellular state.

### Micropapillae are molecularly distinct from neighbouring sessile cells

The second archetypal high-risk morphology, micropapillary growth, is also morphologically biphasic. In 2D sections, tumour cells in areas of micropapillary growth are either seen on a stromal interface, typically as a sessile monolayer, or they are present in micropapillae, in which case the cells have no stromal contact. As an intermediate form, the filigree pattern has been described^30^, in which narrow stacks of extruded or piled-up epithelial cells are seen. Neither micropapillae nor filigree tufts exhibit fibrovascular cores, distinguishing them from true papillae. For the purposes of this study, we refer to the micropapillary non-stromal interfacing regions as florets (Fig.4A^1^), and the flat stromal-interfacing areas as sessile (Fig.4A^2^). To assess the transcriptomic profiles of these morphologically distinct regions we segmented cores with micropapillary growth patterns into sessile and floret AOIs (Fig.4B). After averaging gene expression across cores containing multiple sessile and/or floret growth patterns from the same patient, we proceeded to analyse floret/sessile growth patterns from 12 patients (22 AOIs: 11 floret, 11 sessile). Differential gene expression between florets and sessile regions highlighted four genes (*BGN, IGFBP7, COL4A1* and *LUM)* significantly downregulated in florets compared to sessile areas (p.adj<0.05), (Fig.4C & D). All four are involved in ECM integrity and cohesion: COL4A1 encodes the basement membrane component collagen IV alpha 1, BGN encodes the collagen fibril assembly mediator biglycan, IGFBP7 encodes insulin-like growth factor-binding protein 7 which stimulates cell adhesion and binds collagen IV, and LUM encodes leucine-rich proteoglycan which binds collagen fibrils.

**Figure 4.**
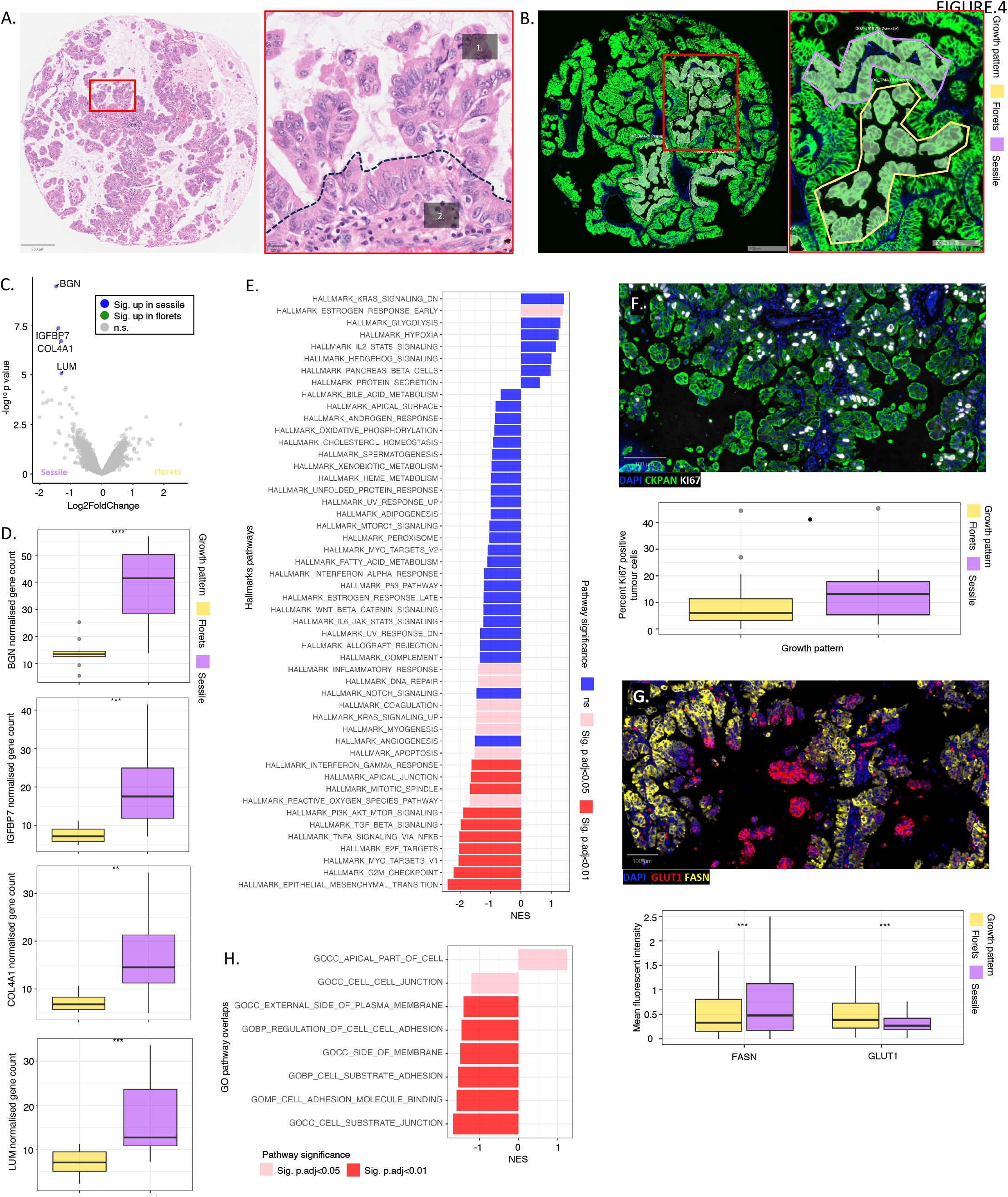
A: Haematoxylin and eosin stained representative image of micropapillary growth pattern (left: scale bar 200μm, right: scale bar 20μm). B: Example segment selection on Digital Spatial Profiler (Nanostring) using manual polygon selection of floret (yellow) and sessile (lilac) regions (left: scale bar 200μm, right: scale bar 100μm). C: Volcano plot of significantly differentially expressed genes (DEG) (p.adj ≤ 0.05). D: Top to bottom: boxplot of target counts in floret and sessile segments for significantly DEGs, *BGN, IGFBP7, COL4A1* and *LUM*. (Mann-Whitney test). E: Gene set enrichment analysis (GSEA) of micropapillary_florets v. micropapillary_sessile segment gene expression using Hallmarks gene-sets (blue: not significant, pink: p.adj ≤0.05, red: p.adj ≤0.01). F: Representative multiplex immunofluorescent image (blue: DAPI, green: CKPAN, white: Ki67) (scale bar 100μm). Boxplot shows percent Ki67 positive tumour cells in floret and sessile segments. G: Representative multiplex immunofluorescent image below (blue: DAPI, red: GLUT1, yellow: FASN) (scale bar 100μm). Boxplot shows mean FASN and GLUT1 fluorescent intensity in floret and sessile segments (Mann-Whitney test). (· p-value < 0.1, * p-value < 0.05, ** p-value < 0.01, *** p-value < 1e-3).

We next performed GSEA using the Hallmarks gene sets (Fig.4E) through which we identified 17 pathways significantly enriched (p.adj≤0.05) in sessile compared to floret regions. These included several proliferation-related pathways, EMT, apoptosis and apical junction components. Immunofluorescence for Ki67 qualitatively confirms that micropapillae are often much less proliferative than their sessile neighbours (Fig.4F) although nominal significance is not reached overall (p=0.095). We also observed a clear increase in pan-cytokeratin fluorescence intensity in micropapillary florets compared to sessile regions as determined by qIF (p=<2.2^e-16^) providing further evidence of a more epithelial/less mesenchymal state for cells in micropapillary florets compared to sessile regions. A metabolic shift between sessile areas and florets (elevated ROS hallmark constituents and a trend towards reduced glycolysis and hypoxia) was also identified by GSEA; we investigated this by quantification of glycolytic and fatty acid metabolism markers with multiplex immunofluorescence. We observed a spatially distinct expression of GLUT1 and FASN immunolabelling with high GLUT1 expression (red) localised to florets (comparison of GLUT1 mean intensity, florets v sessile: p=<0.001) and high FASN expression (yellow) to sessile regions (comparison of FASN mean fluorescence intensity, florets v sessile: p=<0.001) (Fig.4G).

We wished to further investigate the observation that apical junction components were enriched in sessile ROIs. We examined this pathway in more detail through computation of overlapping genes with the gene ontology pathway database (Fig.4H). This analysis highlighted significant enrichment of several pathways related to cell-substrate interactions (“cell substrate junction”, “cell adhesion molecule binding”, “cell substrate adhesion”) and cell-cell interactions (“regulation of cell-cell adhesion”, “cell-cell junction”, “side of membrane”).

Taken together our data highlights several differences between sessile cells and micropapillary cells at both mRNA and protein expression levels and suggests a dynamic model to explain the origin of this clinicopathologically highly significant growth pattern (Fig.5) and explain why some 2D growth patterns become adorned by micropapillae while others do not. The 2D sessile layer component is relatively highly proliferative, with a large fraction of cells in-cycle (Fig.5, panel 1), causing cellular crowding and compression (Fig.5, panel 2). This corresponds to the ‘filigree’ appearances previously described. Ultimately this overcrowding leads to apical extrusion of a cell with separation from the basement membrane, which effectively nucleates the separation of its neighbours from the stroma to accommodate the ongoing overpopulation of the sessile layer (Fig.5, panel 3). Crucially, these extruded cells neither separate from their neighbours, indicative of highly functional apical junctions, and nor do they succumb to anoikis on separation from the basement membrane. Rather, they enter a hyper-epithelial but non-proliferative ‘dormant’ state. This condition is accompanied by, and may rely upon, a hypoxic metabolic switch due to physical separation from their vascular supply. The result is that as cells are displaced from the surface due to crowding, they form chains or tubes of cells linked by intact apical junctions, in a quiescent, hypoxic state (Fig.5, panel 4).

**Figure 5.**
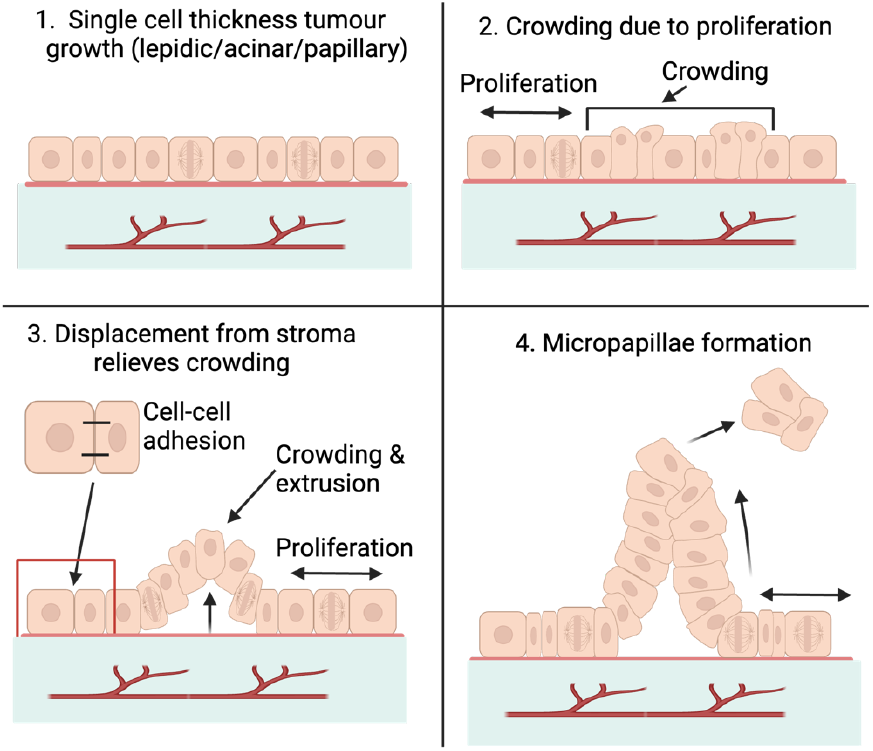
Proposed mechanism of micropapillae development. The 2D sessile layer component is relatively highly proliferative, with a large fraction of cells in-cycle (panel 1), causing cellular crowding and compression (panel 2). This overcrowding leads to apical extrusion of a cell with separation from the basement membrane, which effectively nucleates the separation of its neighbours from the stroma to accommodate the ongoing overpopulation of the sessile layer (panel 3). These extruded cells enter a hyper-epithelial but non-proliferative ‘dormant’ state. This condition is accompanied by, and may rely upon, a hypoxic metabolic switch due to physical separation from their vascular supply. The result is that as cells are displaced from the surface due to crowding, they form chains or tubes of cells linked by intact apical junctions, in a quiescent, hypoxic state (panel 4).

## Discussion

Previous studies of growth patterns of lung adenocarcinoma at the transcriptomic level are few, and due to limitations in the technologies applied generally fail to examine morphologically pure regions^5^. To address this, we applied spatial transcriptomics (Nanostring, Digital Spatial Profiler) for the assessment of growth pattern specific gene expression using the whole transcriptome atlas (18,000 protein coding genes) on formalin fixed paraffin embedded tissue microarrays. Using this approach, we were able to accurately profile gene expression of the full range of WHO-recognised growth patterns and more specifically examine intra-growth pattern morphological heterogeneity in the two best characterised high-risk patterns, namely solid and micropapillary growth.

Initial examination of the data using principal component analysis and hierarchical clustering of GSVA highlighted that while each growth pattern exhibited distinct biological hallmarks, 3 clusters of growth patterns were resolved based on hallmark commonalities. Interestingly, solid and micropapillary growth patterns were associated with different groups, suggesting different molecular origins and biological mechanisms. Furthermore, a pseudo-time analysis identified two distinct trajectories, both starting at normal alveolar epithelial gene expression, sharing a path through low- and medium-grade appearances, and then diverging sharply into solid and micropapillary growth. This ordering both recapitulates previous findings about growth pattern transition from low to high-grade^31^ and suggests a point of molecular divergence between solid and micropapillary patterns. The position on the trajectory demonstrates potential clinical relevance of the identified signatures, as shown by association studies with OS.

Taken together we observe that the low- and medium-grade growth patterns (in situ, papillary and acinar tumour regions) are all essentially 2-dimensional in their epithelial components and consist of sheets of malignant cells growing on minimally changed interstitium, or on an arborising vascular tree, or as malignant glands in desmoplastic stroma, respectively. In contrast to this, the high-grade patterns, solid, micropapillary, and cribriform, represent a switch to 3-dimensional arrangements of tumour cells. Within this comparison, the two archetypal high-risk growth patterns, micropapillae and solid tumour islands, represent 2 highly divergent ways in which malignant cells that can no longer be accommodated in a 2-dimensional sheet anchored to stroma may survive. In solid growth, the cells have undergone a degree of dedifferentiation, polarity loss, and EMT, and are able to grow as a solid mass with a highly proliferative and stromally invasive periphery. Micropapillae also enable tumour cells to extend away from the 2D stromal interface, but by a wholly different mechanism. Our evidence supports a dynamic model of ‘upward’ extrusion of cells and maintained or even accentuated epithelial identity and polarity within the micropapillae. The solid programme depends upon a heavy transcriptional drive through E2F and MYC and the ongoing invasiveness of the periphery; the micropapillary programme is the consequence of local epithelial dynamics as outlined in our proposed dynamic model.

We suggest, therefore, that there may often be a true evolutionary progression from in situ growth through an intermediate 2-dimensional morphology. As increasingly proliferative 2-dimensional clones emerge, selection becomes more acute as cells die due to overcrowding and competition, leading ultimately to selection of a new survival strategy. This can be ‘upward’ extrusion and micropapilla formation, or ‘downward’ invasion, EMT, and solid growth. These two highly divergent strategies are equally lethal, as assessed by the hazard ratios associated with their fractional presence in lung adenocarcinomas. They are both highly metastatic, but we suggest that the underlying mechanisms of metastatic progression may differ. We show that the peripheral elements of solid islands of primary human lung adenocarcinoma exhibit proliferative and motile programs, consistent with their highly destructive behaviour, and it is likely that they achieve distant metastasis by direct infiltration of vascular structures. Micropapillae have different properties; they are metabolically indolent and hyper-epithelial but originate plastically from relatively proliferative and EMT-transitioned sessile cells. We suggest that they are the ideal metastatic agent, ready to survive the rigors of vascular transit with little to no requirement for nutrition or oxygen, but poised to re-enter a proliferative state when they encounter a suitable niche. How they enter the vascular system is less clear; we suggest that they fragment and subsequently obtain vascular ingress, perhaps by passive lymphatic suction into leaky or abnormal terminal vessels.

These divergent models have implications for systemic therapy. For example, cytotoxic therapies targeting proliferative cells may be effective on solid island peripheries, killing the population of cells most likely to immediately invade and metastasise. In addition, our finding of the centrality of the E2F family of transcription factors suggests that this morphology may be susceptible to E2F family inhibition. But micropapillae, being less proliferative and relatively metabolically inactive, are likely to be more resistant to such therapies, in addition to being harder to reach with agents delivered via the circulatory system. They might even be displaced or released by therapies which kill the sessile component or cause fragmentation of micropapillae.

Our study is limited to an examination of the epithelial components of these growth patterns; there are undoubtedly additional phenotypes of the fibroblastic components related to growth pattern, and these are currently under study. In addition, we cannot address interesting questions about differences in the immune microenvironment. Finally, we do not address the issue of complex glandular growth in detail, the complexity of which lies beyond the scope of this focussed study. However, we note the colocalization of acinar, solid and cribriform patterns in our PCA analysis, supporting the idea that these morphologies lie on a molecular continuum.

In conclusion, through our targeted spatially resolved transcriptomic assessment of non-small cell lung adenocarcinoma growth patterns we identify distinct differences between growth patterns and uncover evidence that solid and micropapillary patterns may arise through different evolutionary trajectories. These two highly lethal patterns are both shown to have marked plasticity related to hypoxic status, and the hypoxic fraction of micropapillary areas appears to bear the most immediate clinical hazard through a mechanism of passive quiescence, making them the ideal agent for metastatic spread. Consideration of high-risk growth patterns as biologically distinct entities may yield novel therapeutic options for patients presenting with these highly divergent forms of malignancy.

## Acknowledgments

The authors gratefully acknowledge the support of NHS Research Scotland (NRS) Greater Glasgow and Clyde Biorepository.

## Data availability statement

The data generated in this study are available upon request from the corresponding author.

## Figure legends

**Figure S1.**
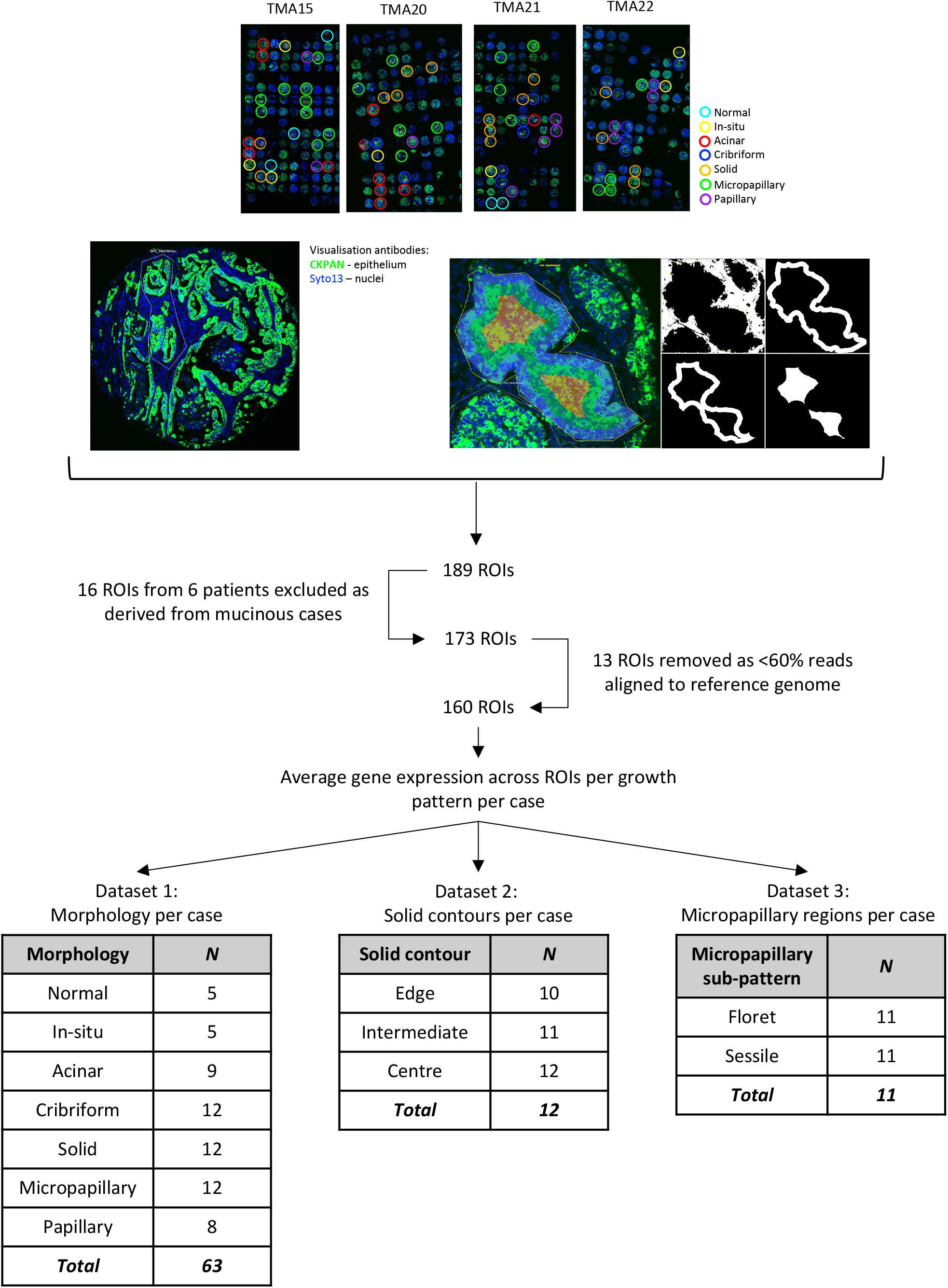
Summary of experimental workflow. Top to bottom: Four tissue microarrays comprising cores representing a range of primary lung adenocarcinoma morphologies. Sampled cores circled (colour key: cyan – normal, yellow – in-situ, red – acinar, blue – cribriform, orange – solid, green – micropapillary, purple – papillary). Primary visualisation markers (CKPAN = epithelium, Syto13 = DNA/nuclei) were applied to facilitate segment selection. A variety of segment selection strategies were employed including manual polygon selection and “contouring” (example image shown). In total, 189 regions of interest (ROIs) were sampled using the Digital Spatial Profiler (DSP, Nanostring). Sixteen ROIs were removed as these samples originated from 6 patients with mucinous histology. Following pre-analytical quality control steps (including exclusion of segments with <60% aligned reads), 160 segments were used for downstream analysis and organised into three datasets for further analysis (Dataset 1: morphology by case, raw counts from segments of the same morphology from the same patient were averaged deriving a case_morph sample (*n=63)*. Dataset 2: three solid contours were taken from the tumour/stroma interface to the centre of solid “islands”. Multiple individual contours (e.g., edge) from the same patient were averaged deriving a contour per case sample (*n=12)*. Dataset 3: floret and sessile segments were derived from cores of micropapillary growth patterns. Multiple floret or sessile segments from the same patient were averaged deriving floret and sessile samples (*n=11)*.

**Figure S2.**
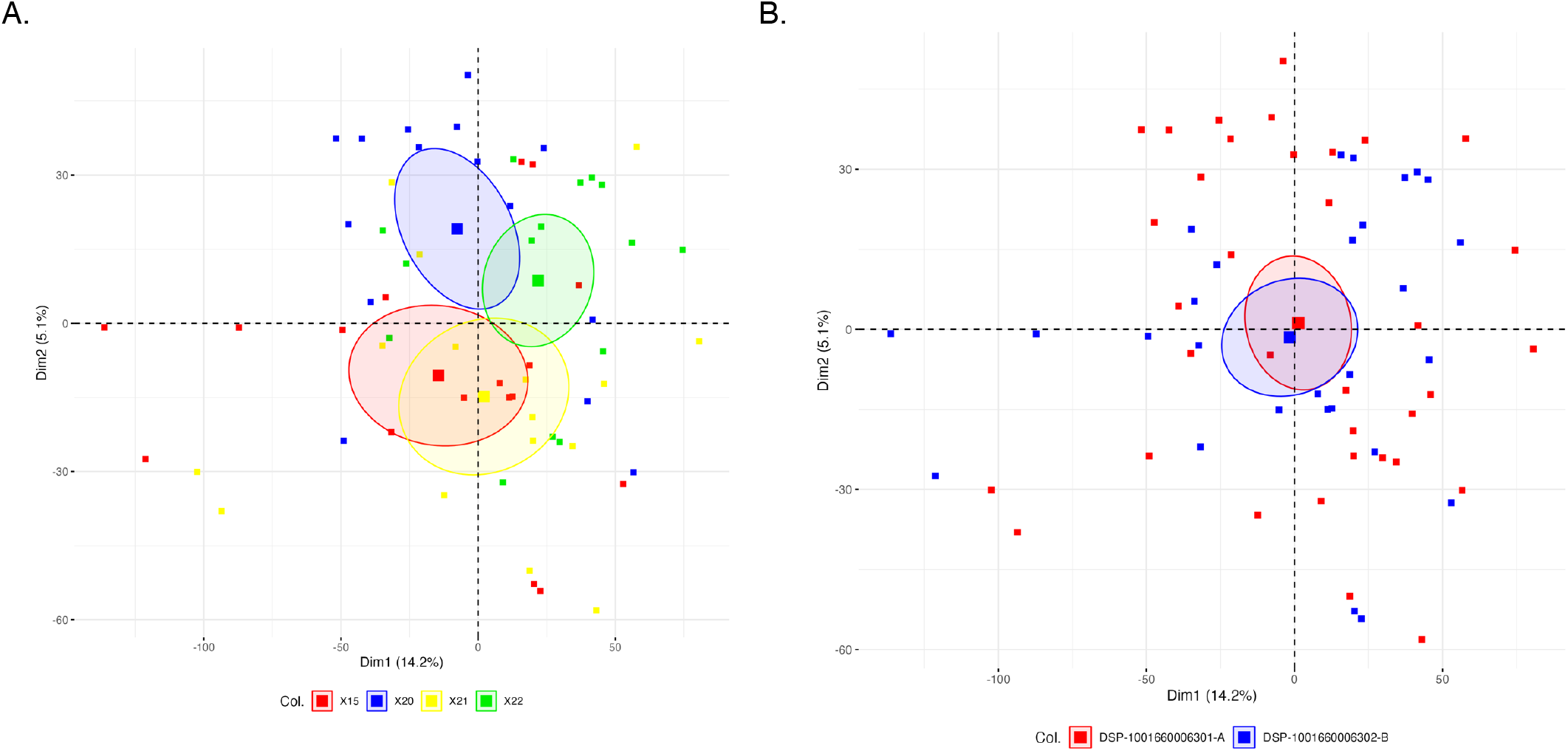
Batch effect assessment. A: Principal component analysis plot of averaged gene expression from dataset 1 coloured by tissue microarray block (red: TMA15, blue: TMA20, yellow: TMA21, green: TMA22). B: Principal component analysis plot of averaged gene expression from dataset 1 coloured by sequencing plate (red: A, blue: B).

**FigureS2.**
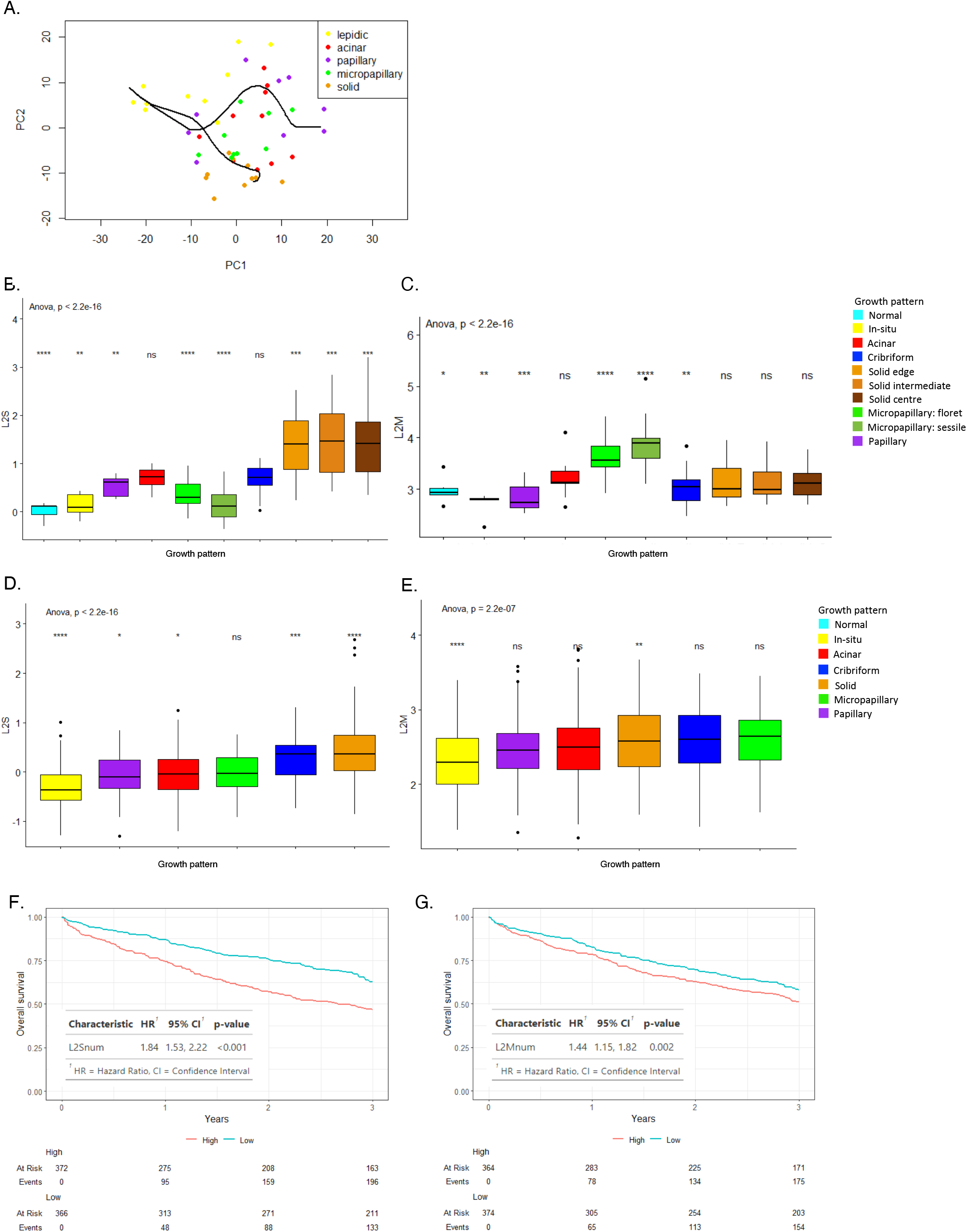
A: Boxplot of L2S score in function of histological pattern in pure epithelial samples. P-values are the result of t-test between corresponding histological pattern and all the others. B: Boxplot of L2M score in function of histological pattern in pure epithelial samples. C: Boxplot of L2S score in function of histological pattern in bulk samples. D: Boxplot of L2M score in function of histological pattern in bulk samples. E: Survival analysis of L2S. Top panel is the Kaplan-Meier representation, L2S is discretised in low and high based on median value. Bottom panel represents corresponding table of estimates of corresponding Cox regression analysis. F: Survival analysis of L2M. Top panel is the Kaplan-Meier representation, L2M is discretised in low and high based on median value. Bottom panel represents corresponding table of estimates of corresponding Cox regression analysis. (· p-value < 0.1, * p-value < 0.05, ** p-value < 0.01, *** p-value < 1e-3).

